# Lysosomal vesiculophagy terminates STING signalling

**DOI:** 10.1101/2022.03.20.485012

**Authors:** Yoshihiko Kuchitsu, Kojiro Mukai, Yuki Takaada, Ayumi Shinojima, Shiori Hamano, Rei Uematsu, Emari Ogawa, Takehiro Suzuki, Naoshi Dohmae, Takefumi Uemura, Hiroyuki Arai, Satoshi Waguri, Tomohiko Taguchi

## Abstract

Stimulator of interferon genes (STING) is essential for the type I interferon response against a variety of DNA pathogens^1,2^. Upon emergence of cytosolic DNA, STING translocates from the endoplasmic reticulum (ER) to the Golgi where STING activates the downstream kinase TBK1, then to lysosome through recycling endosomes (REs) for its degradation^3–6^. Although the molecular machinery of STING activation is extensively studied and defined^7^, the one underlying STING degradation has not yet been fully elucidated. Here we show an unanticipated mechanism termed “lysosomal vesiculophagy” dictates STING degradation. Airyscan super-resolution microscopy and correlative light/electron microscopy show that a cluster of STING-positive vesicles of an RE origin are directly encapsulated into lysosome. Screening of mammalian *Vps* genes, the yeast homologues of which regulate Golgi-to-vacuole transport^8^, shows that the endosomal sorting complexes required for transport (ESCRT) is essential for the STING encapsulation into lysosome. Knockdown of Tsg101 and Vps4, components of ESCRT^9^, results in the accumulation of STING vesicles in the cytosol, leading to the sustained type I interferon response. STING undergoes ubiquitination at REs and degradation of STING requires the ubiquitin-binding domain of Tsg101, ensuring the selective degradation of activated STING. Our results reveal a novel mode of autophagy that prevents hyperactivation of innate immune signalling and provide insights into the ER-Golgi-REs-lysosomes pathway for degradation of transmembrane proteins.

---

Lysosomes are membrane-bound organelles and contain various acid hydrolases to degrade macromolecules including proteins, lipids, and nucleotides^10,11^. The degradative substrates in the extracellular space or at the plasma membrane are delivered to lysosomes by the endocytic pathway, whereas the ones in the cytosol reach lysosomes by a mechanism designated autophagy^12^. There exist at least three distinct types of autophagy, *i.e*., macroautophagy (delivery of cytosolic substrates to lysosomal lumen via autophagosomes)^13^, chaperone-mediated autophagy (CMA: translocation of cytosolic substrates to lysosomal lumen directly across the limiting membrane of lysosomes)^14^, and microautophagy (an inward invagination of the limiting membrane of lysosomes, generating intraluminal vesicles destined for degradation)^15^. Mechanism and biological significance of macroautophagy and CMA have been extensively investigated, in contrast, those of microautophagy remain unclear, in particular, in mammalian cells^16^.

STING is an ER-localized transmembrane protein essential for control of infections of DNA viruses and tumor immune surveillance^2^. After binding to cyclic GMP-AMP (cGAMP)^17^, which is generated by cGAMP synthase (cGAS)^18^ in the presence of cytosolic DNA, STING translocates to the Golgi where STING recruits TBK1 from the cytosol and triggers the type I interferon- and proinflammatory responses through the activation of interferon regulatory factor 3 (IRF3) and nuclear factor-kappa B (NF-kB)^3,4,6,19,20^. STING further translocates to lysosomes for its degradation through REs, so that the STING-triggered immune signalling is terminated^5,6,21–25^. The mechanism of how STING, a transmembrane protein on the exocytic membrane traffic, reaches lysosomes has been poorly understood.

## Direct encapsulation of STING into lysosomal lumen

To examine how STING is delivered to lysosome, *Sting^-/-^* mouse embryonic fibroblasts (MEFs) were stably transduced with mRuby3-tagged mouse STING and EGFP-tagged Lamp1 (a lysosomal protein), and imaged with Airyscan super-resolution microscopy. Without stimulation, mRuby3-STING localized to a reticular network throughout the cytoplasm (Fig. 1a), suggesting that STING localized at ER^3^. mRuby3-STING diminished 12 hours after stimulation with DMXAA (a membrane-permeable STING agonist). In contrast, addition of lysosomal protease inhibitors (E64d/pepstatin A) restored the fluorescence, with mRuby3-STING mostly in lysosomal lumen, not at the limiting membrane of lysosome (Fig. 1b). These results suggested that degradation of STING proceeded in lysosomal lumen.

**Fig. 1.**
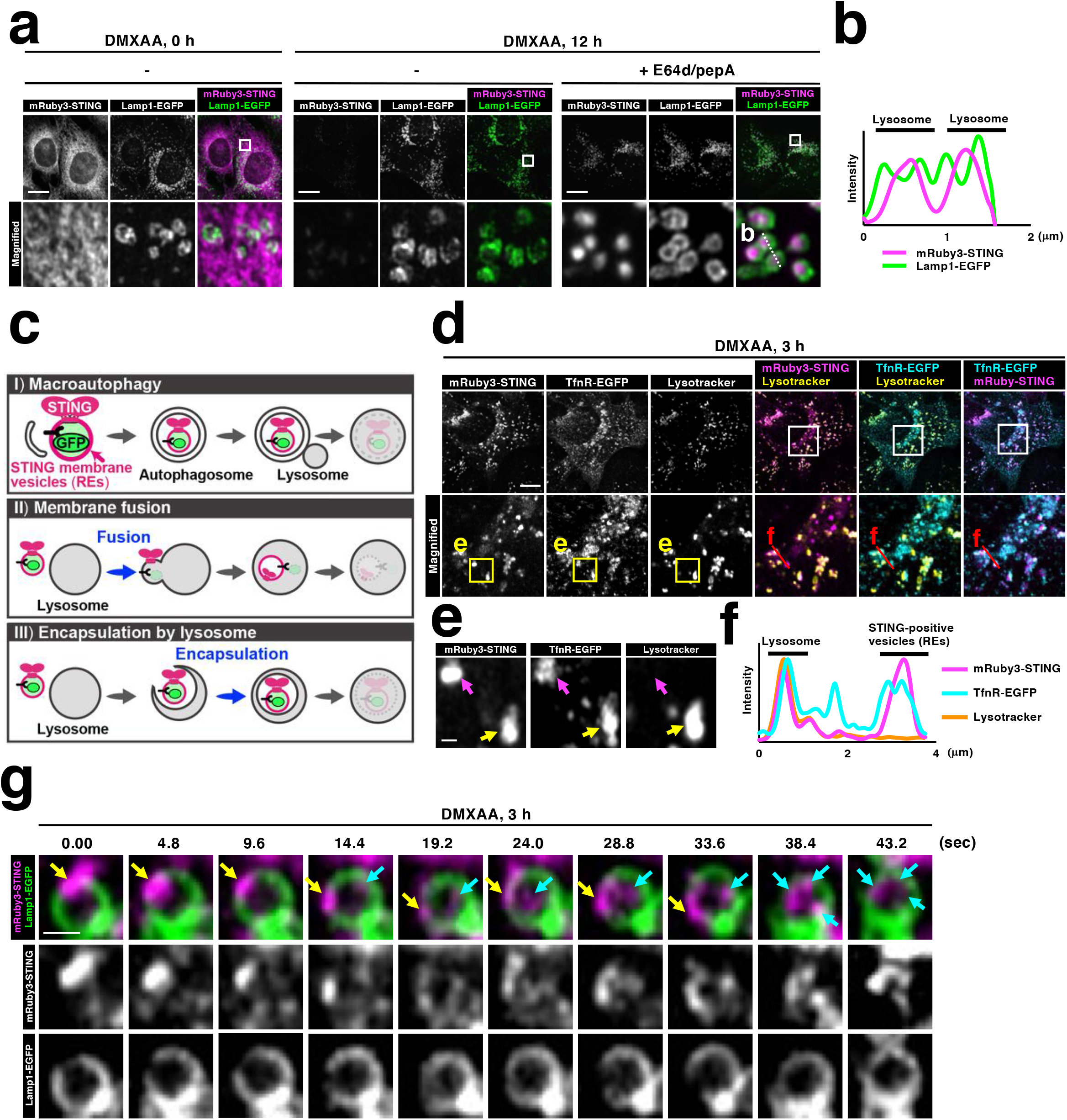
Direct encapsulation of STING into lysosomal lumen. **a**, *Sting^-/-^* MEFs stably expressing mRuby3-STING (magenta) and Lamp1-EGFP (green) were treated with DMXAA (25 μg ml^-1^) for 0 or 12 h. For the inhibition of lysosomal proteolysis, E64d (30 μg ml^-1^) and pepstatin A (40 μg ml^-1^) were added to the medium. Cells were fixed and imaged by Airyscan super-resolution microscopy. The boxed areas in the top panels are magnified in the bottom panels. Scale bars, 10 μm. **b**, Fluorescence intensity profile along the white dotted line in (**a**) is shown. **c**, Three ways to deliver STING to lysosomal lumen: (I) “Macroautophagy”; STING vesicles are first occluded into autophagosomes, which then fuse with lysosome. (II) “Membrane fusion”; STING vesicles fuse with lysosome, followed by invagination of limiting membrane of lysosome, yielding intraluminal STING vesicles. (III) “Encapsulation by lysosome”; STING vesicles are directly encapsulated into lysosomes. **d**, TfnR-EGFP (cyan) and mRuby3-STING (magenta) were stably expressed in *Sting^-/-^* MEFs. Cells were treated with DMXAA for 3 h and then with LysoTracker Deep Red (yellow). Live cell imaging was performed with Airyscan superresolution microscopy. The boxed areas in the top panels are magnified in the bottom panels. Scale bar, 10 μm. **e**, The yellow boxed areas in (**d**) are magnified. The yellow arrow indicates lysosome positive with mRuby3-STING and TfnR-EGFP. The magenta arrow indicates REs, in which mRuby3-STING co-localized with TfnR-EGFP. Scale bar, 500 nm. **f**, Fluorescence intensity profile along the red line in (**d**) is shown. **g**, *Sting^-/-^* MEFs stably expressing mRuby3-STING and Lamp1-EGFP were imaged by Airyscan super-resolution microscopy every 0.4 seconds from 3 hours after DMXAA stimulation. Selected frames from Supplementary Video 2 are shown (see also Extended Data Fig.3a and 3b). The yellow arrows indicate a cytosolic STING chunk in close proximity to lysosomal limiting membrane. The cyan arrows indicate STING inside lysosome. Scale bar, 500 nm.

Membrane proteins, such as STING, may have access to lysosomal lumen by three ways, *i.e*., (i) macroautophagy, (ii) membrane fusion, or (iii) direct encapsulation (Fig. 1c). Several reports suggested that STING degradation did not require macroautophagy^21,23,25^, and we confirmed this in Atg5 tet-off MEFs in which macroautophagy was impaired in the presence of doxycycline^26^ (Extended Data Fig. 1a). Importantly, with lysosomal protease inhibitors, mRuby3-STING accumulated in lysosomal lumen in Atg5-depleted cells (Extended Data Fig. 1b), suggesting that macroautophagy was not involved in the delivery of STING into lysosomal lumen. The other two scenarios can be distinguished by probing the luminal pH of STING vesicles. We exploited an RE protein transferrin receptor (TfnR). TfnR was *C*-terminally tagged with EGFP, which thus faced the lumen of REs. If “membrane fusion” occurs, the fluorescence of EGFP should be quenched because of its exposure to lysosomal acidic milieu^27^. If “direct encapsulation” occurs, the fluorescence of EGFP should linger until two membranes surrounding EGFP are digested by lysosomal lipases. TfnR-EGFP was expressed together with mRuby3-STING. mRuby3-STING started to co-localize with TfnR-EGFP 60 min after DMXAA stimulation (Extended Data Fig. 2, Supplementary Video 1), showing that STING reached REs by that time^3,6^. Intriguingly, the fluorescence of TfnR-EGFP was detected at lysosomes (lysotracker: a lysosomal indicator) together with mRuby3-STING 3 hours after DMXAA stimulation (Fig. 1d-f). These results suggested that the STING delivery to lysosomal lumen was mediated by “direct encapsulation”.

We then performed time-lapse imaging of live cells. Cells were imaged every 0.4 seconds from 3 hours after DMXAA stimulation. We often found that an irregularly shaped mRuby3-STING-positive chunk in close proximity to lysosomal limiting membrane, translocated into lysosomal lumen (Fig. 1g, Extended Data Fig. 3, Supplementary Video 2). During this process, mRuby3-STING appeared not to diffuse along lysosomal limiting membrane, supporting the mechanism of “direct encapsulation”. mRuby3-STING did not co-localize with Rab5-positive early endosomes up to 4 hours after DMXAA stimulation (Extended Data Fig. 1c-h), suggesting that the delivery of STING to lysosomes was not via early endosomes.

## An evidence of “direct encapsulation” by CLEM

We sought to validate “direct encapsulation” by another approach. “Direct encapsulation”, but not “membrane fusion”, will result in the generation of a limiting membrane (moss green in Fig. 2d) that surrounds STING vesicles. To examine whether STING vesicles in lysosomal lumen is surrounded by membrane, correlative light and electron microscopy (CLEM) was exploited. Cells were fixed and imaged with Airyscan super-resolution microscopy 3 hours after DMXAA stimulation. The same cells were then processed for electron microscopy. Two images in the same region of cells from fluorescence microscopy and electron microscopy were aligned according to multiple lysosomal positions (Fig. 2a). The CLEM analysis revealed that a STING-positive chunk inside lysosomes (magenta in Fig. 2b, c, Extended Data Fig. 4a, b) was composed of a cluster of membrane vesicles. Importantly, the cluster of membrane vesicles was surrounded by single membrane (indicated by arrows in moss green), demonstrating that “direct encapsulation” is a mechanism underlying the STING delivery into lysosomal lumen. The CLEM analysis also showed the nature of STING membranes that were free from or associated with lysosomes (Fig. 2e, f, Extended Data Fig. 4c, d). These irregularly shaped STING-positive chunks were indeed clusters of vesicles with electron-dense coat (60 - 130 nm in diameter) (Fig. 2h). Together with the data of live cell imaging (Fig. 1d, g), these results suggested that a cluster of vesicles with an RE origin was directly encapsulated into lysosomes. We thus coined this process “lysosomal vesiculophagy”.

**Fig. 2.**
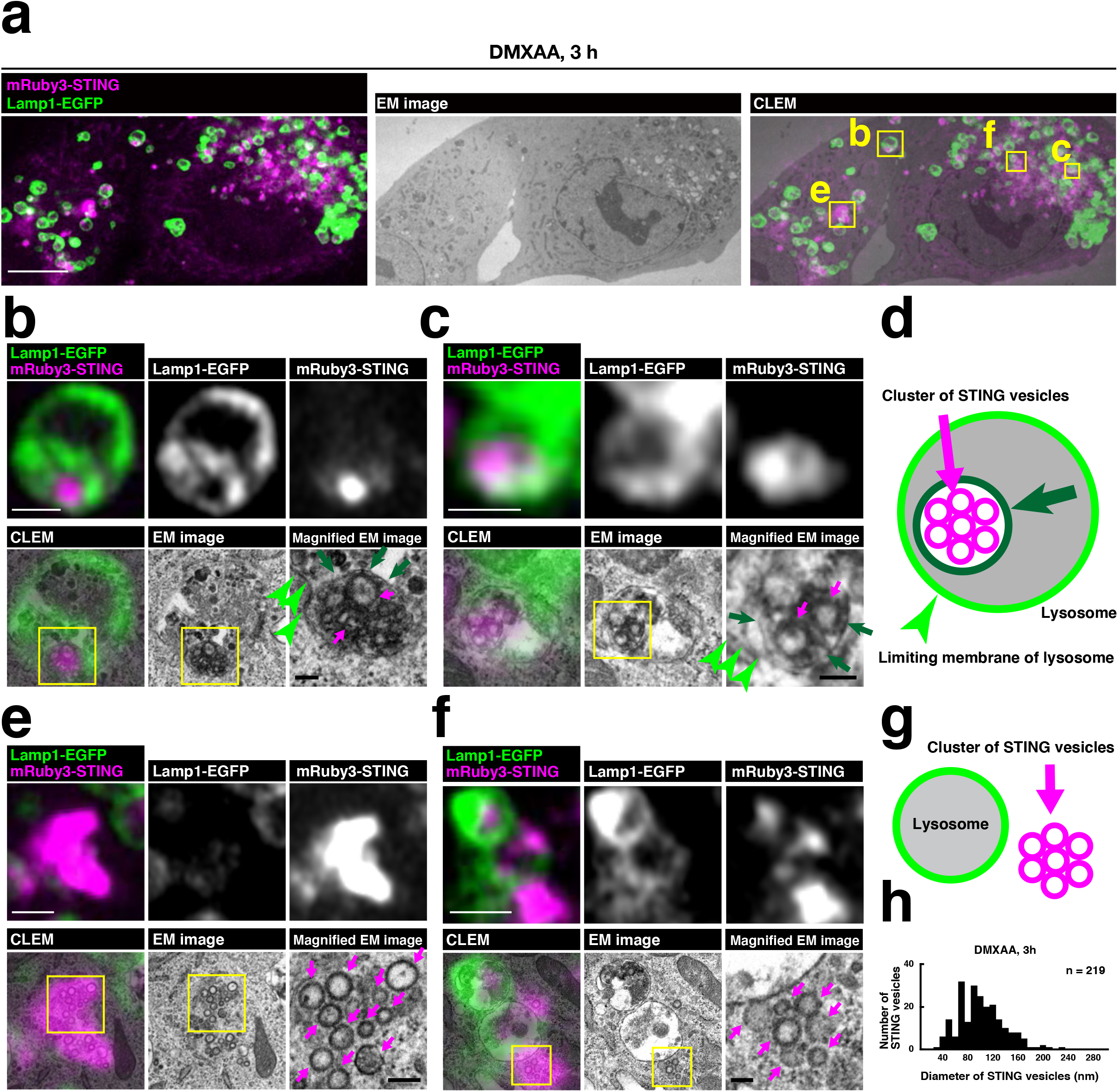
An evidence of “direct encapsulation” by CLEM. **a-h**, *Sting^-/-^* MEFs stably expressing mRuby3-STING (magenta) and Lamp1-EGFP (green) were treated with DMXAA (25 μg ml^-1^) for 3 h in the presence of E64d/pepstatin A/orlistat (lipase inhibitor). Lamp1-positive lysosomes and STING-positive vesicles (or structures) were identified by Airyscan super-resolution microscopy before processing for transmission electron microscopy to examine their ultrastructure (**a**). The boxed areas in (**a**) are magnified in (**b**, **c**, **e**, and **f**). For EM images of serial sections, see Extended Data Fig. 4. A graphical image of lysosome containing membrane-encapsulated STING vesicles (**d**). Arrows in moss green in (**b**, **c**, and **d**) indicate the membrane that surrounds STING vesicles. Arrowheads in light green indicate limiting membrane of lysosome. A graphical image of the lysosome and cytosolic STING vesicles not encapsulated by membranes. (**g**). The diameter of STING-positive membrane vesicles was measured and plotted as histograms (**h**). Scale bars, 10 μm in (**a**), 500 nm in (**b**, **c**, **e**, and **f**), 100 nm in magnified EM image

## STING degradation and termination of type I interferon response require ESCRT proteins

In yeast, more than 40 proteins were designated vacuolar protein sorting (Vps) proteins^8,28^, which function in sorting of newly synthesized vacuolar proteins from late-Golgi to vacuole (the yeast equivalent of lysosome). Given the analogous traffikcing pathways that STING and vacuolar proteins follow, we reasoned if mammalian Vps homologues regulate STING traffic to lysosomes. The impaired traffic of STING to lysosomes should result in the suppression of STING degradation^5,6,21–25^, and may also in the duration of the STING-triggered type I interferon response.

We screened 75 *Vps* mammalian homologues with siRNAs in two criteria, *i.e*., the effect on STING degradation and termination of the type I interferon response. The degree of degradation and that of the type I interferon response were quantitated using flow cytometer and type I interferon bioassay, respectively (Fig. 3a). Knockdown of 55 *Vps* genes showed enhanced suppression of STING degradation, compared to that with control siRNA (Fig. 3b, Extended Data Fig. 5a). Atp6v1b2, a component of subunit B of v-ATPase, was included in this assay as a positive control. Knockdown of 40 *Vps* genes showed an increased type I interferon response, compared to that with control siRNA (Fig. 3c, Extended Data Fig. 5b). The genes that were ranked within top 25 in both criteria were selected and listed (Fig. 3d). These genes included 4 *Vps* genes (*Vps28, Tsg101, Vps37a*, and *Chmp4b*) that belong to ESCRT, *Vps4* (AAA-ATPase), and *Vps39* [a subunit of homotypic fusion and vacuole protein sorting (HOPS) complex]. Knockdown of these genes significantly enhanced the expression of Cxcl10, a STING-downstream gene, compared to that with control siRNA (Fig. 3e, Extended Data Fig. 5c), corroborating the results with the type I interferon bioassay (Fig. 3c).

**Fig. 3.**
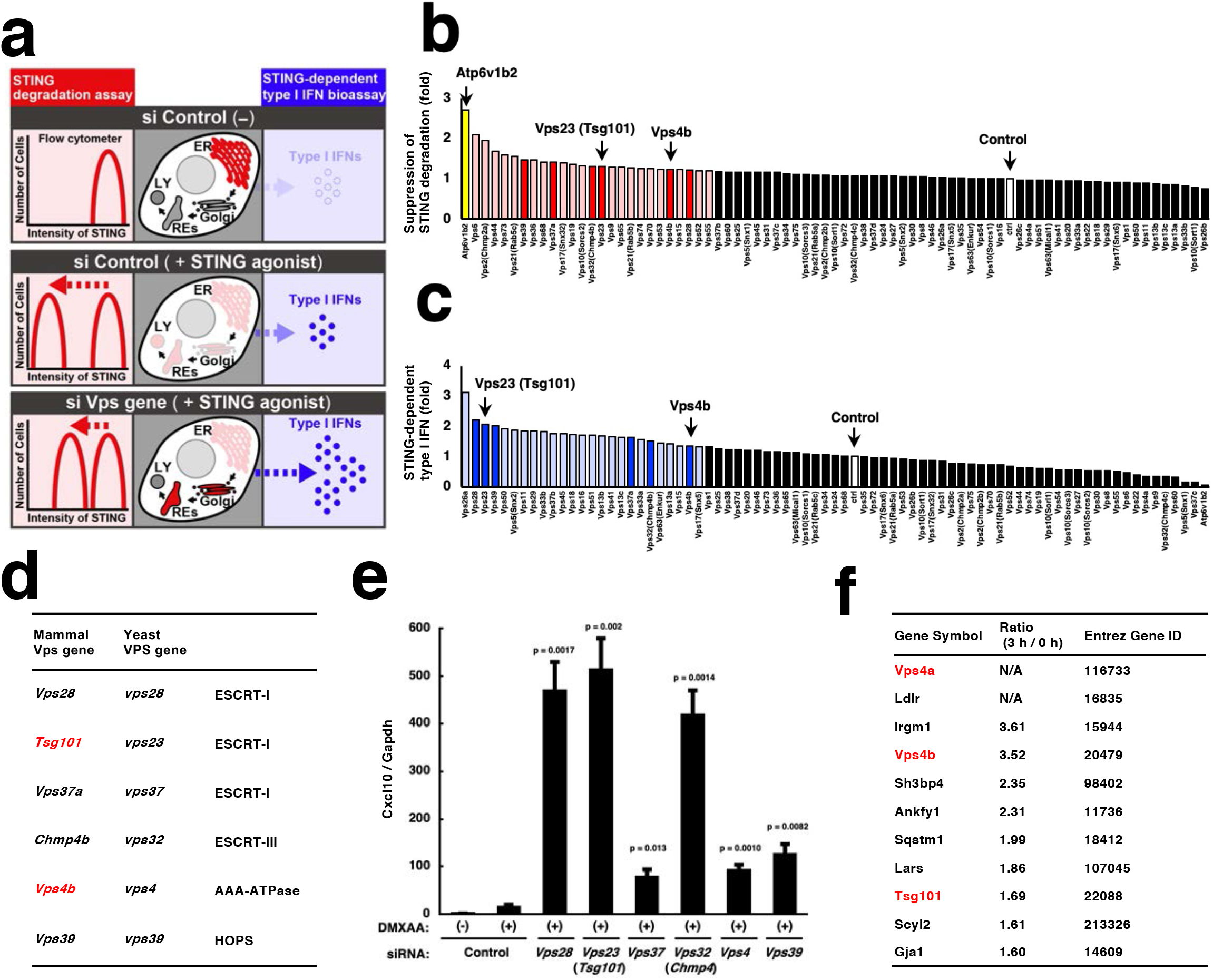
Screening of mammalin *Vps* genes that are essential for STING degradation and termination of type I interferon response. **a**, A schematic overview of the screening procedures. **b**, Screening of mammalian *Vps* genes required for degradation of STING. *Sting^-/-^* MEFs reconstituted with mRuby3-STING were treated with siRNA for the indicated *Vps* genes for 54 h, and stimulated with DMXAA for 18 h. Cells were fixed and analyzed by flow cytometry. Mean fluorescence intensity (MFI) of mRuby3 in stimulated cells was divided by MFI of mRuby3 in the correponding unstimulated cells. The calculated value from cells treated with *Vps* siRNA was then normalized to that of cells treated with control siRNA, and the normalized values were listed in decreasing order. The top 25 genes are highlighted in red. Bright red bars indicate the genes that were also ranked within top 25 in (**c**). **c**, Screening for mammalian *Vps* genes required for suppression of STING-dependent type I interferon response. MEFs were treated with siRNA for the indicated *Vps* genes for 62 h, and stimulated with DMXAA for 10 h. Cell supernatants were then collected and analyzed for type I interferon (IFN). Type I interferon activity from cells treated with *Vps* siRNA was normalized to that of cells treated with control siRNA, and the normalized values were listed in decreasing order. The top 25 genes are highlighted in blue. Bright blue bars indicate the genes that were also ranked within top 25 in (**b**). **d**, The *Vps* genes that were ranked within top 25 both in (**b**) and (**c**) are shown. **e**, Quantitative real-time PCR (qRT-PCR) of the expression of Cxcl10 in MEFs that were treated with siRNA for the indicated *Vps* genes for 60 h and then stimulated with DMXAA for 12 h. **f**, FLAG-STING-reconstituted *Sting^-/-^* MEFs were stimulated with DMXAA for 3 h, and lysed. FLAG-STING in the lysates was immunoprecipitated with anti-FLAG M2 antibody. Co-immunoprecipitated proteins were identified by mass spectrometry. The ratio of abundance of identified proteins before and after stimulation was then calculated individually. The listed are lysosomal proteins that showed increased abundance after stimulation. Gene ontology analysis in Uniprot was performed to identify lysosomal proteins. “N/A” indicates a protein that was not detected without stimulation.

We also performed proteomic analysis of FLAG-STING-binding proteins, aiming at identifying proteins that regulate STING degradation at lysosomes. The amount of individual proteins in the immunoprecipitates by anti-FLAG antibody was quantitated before and after stimulation (Supplementary Table 1). We selected the proteins, the amount of which increased after stimulation, and further screened them if they were annotated to “lysosome” in gene ontology in Uniprot. This approach led to identify three Vps proteins (Vps4a, Vps4b, and Tsg101) (Fig. 3f). Together with the aforementioned results (Fig. 3d), we examined the role of Vps4a, Vps4b, and Tsg101 in lysosomal degradation of STING in the subsequent experiments.

## ESCRT proteins function in encapsulation of STING into lysosomal lumen

We sought to identify the site of actions of Tsg101 and Vps4a/4b, thus examined the trafficking of STING in Tsg101- or Vps4a/b-knockdown cells. In cells treated with control siRNA, the fluorescence of mRuby3-STING diminished entirely 12 hours after stimulation with DMXAA, because of its lysosomal degradation (Fig. 4a, b). In contrast, in cells treated with Tsg101 or Vps4a/b siRNA, the fluorescence of mRuby3-STING lingered and co-localized with TfnR, indicating that the transport of STING from REs to lysosomes was impaired. Phosphorylated TBK1, a hallmark of STING activation, also lingered and co-localized with mRuby3-STING in cells treated with Tsg101 or Vps4a/b siRNA (Fig. 4a, b), being consistent with the duration of the STING signalling in these cells (Fig. 3c, e). CLEM analysis of Tsg101- or Vps4a/b siRNA-treated cells showed a cluster of STING-positive vesicles that were peripherally associated with lysosomal limiting membrane (Fig. 4c-j, Extended Data Fig. 6a-d). These results suggested that Tsg101 and Vps4a/4b were essential for encapsulation of a cluster of STING-positive vesicles into lysosomal lumen.

**Fig. 4.**
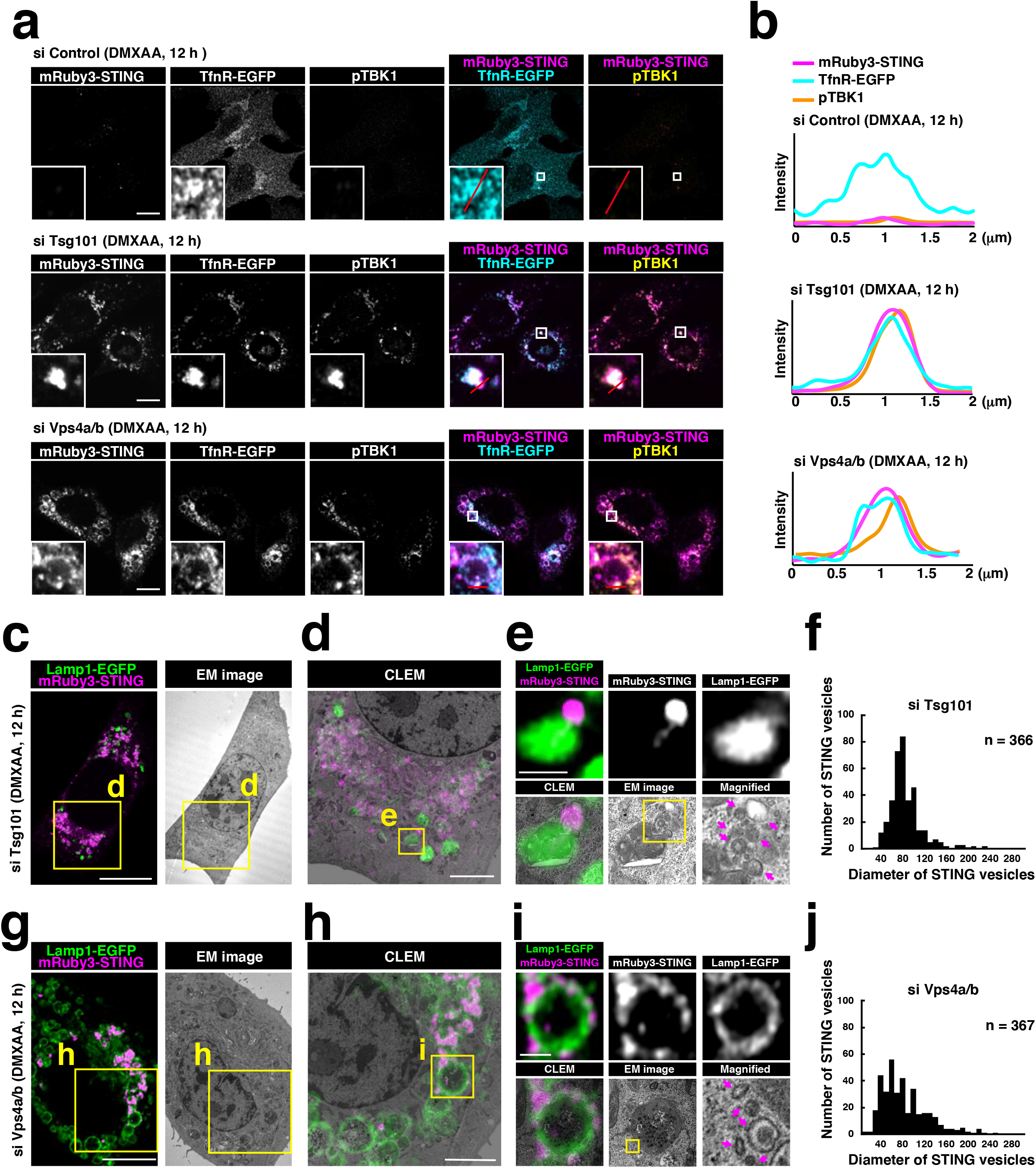
ESCRT proteins function in encapsulation of STING into lysosomal lumen. **a**, TfnR-EGFP (cyan) and mRuby3-STING (magenta) were stably expressed in *Sting^-/-^* MEFs. Cells were treated with the indicated siRNAs for 60 h, and then stimulated with DMXAA for 12 h. Cells were then fixed, permeabilized, and immunostained with anti-pTBK1 (yellow) antibody. Scale bars, 10 μm. **b**, Fluorescence intensity profiles along the red lines in (a) are shown. **c-j**, CLEM analysis of STING-positive vesicles. *Sting^-/-^* MEFs stably expressing mRuby3-STING (magenta) and Lamp1-EGFP (green) were treated with siRNA against *Tsg101* (**c**) or *Vps4a/b* (**g**) for 60 h, and then stimulated with DMXAA for 12 h. Lamp1-positive lysosomes and STINGpositive membranes were identified by Airyscan super-resolution microscopy before processing for transmission electron microscopy to examine their ultrastructure (**c**, **g**). The yellow boxed areas in (**c**) and (**g**) are magnified in (**d**) and (**h**), respectively. The yellow boxed areas in (**d**) and (**h**) are magnified in (**e**) and (**i**), respectively. STING-positive vesicles in (**e**) and (**i**) are indicated by magenta arrows. The diameters of STING-positive vesicles in Tsg101- or Vps4a/b-depleted cells were measured and plotted as histogram (**f**, **j**). Scale bars, 10 μm (**c**, **g**), 2 μm (**d**, **h**), and 500 nm (**e**, **i**).

## Ubiquitin-binding domain of Tsg101 is required for termination of the STING signalling

Given that STING undergoes ubiquitination after stimulation^7^ and Tsg101 binds to ubiquitinated proteins^29,30^, we reasoned that the binding of Tsg101 to ubiquitinated STING would be required for STING degradation and thus termination of the STING signalling. We confirmed the stimulation-dependent ubiquitination of STING by co-immunoprecipitation analysis. STING became extensively ubiquitinated 2 hours after DMXAA stimulation at 37 °C, but not at 20 °C (Fig. 5a). Since low temperature (20 °C) impairs membrane traffic from the trans-Golgi network^31^, these results suggested that the extensive ubiquitination of STING occurs at the post-Golgi compartments.

**Fig. 5.**
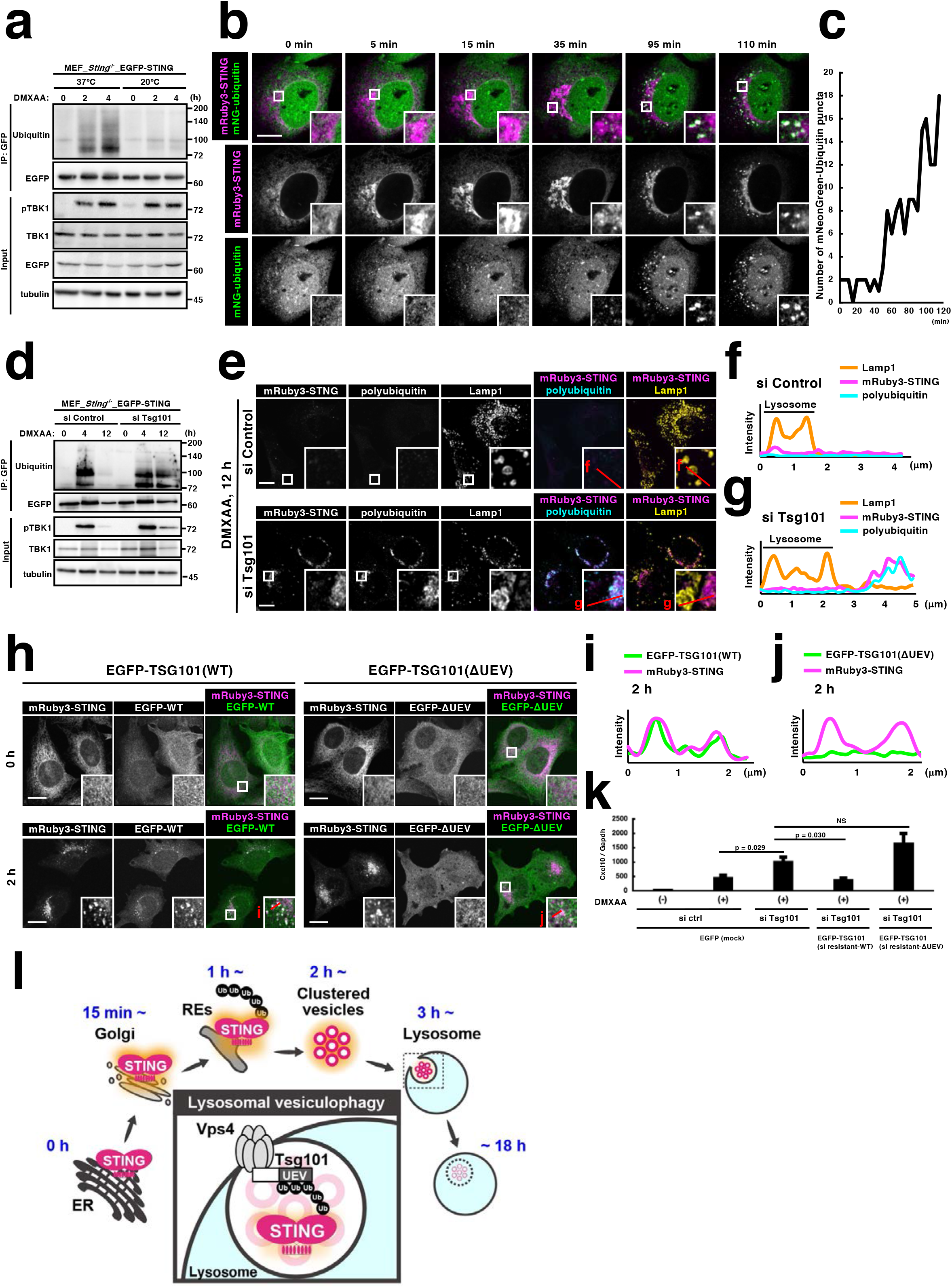
Ubiquitin-binding domain of Tsg101 is required for termination of the STING signalling. **a**, *Sting^-/-^* MEFs reconstituted with EGFP-STING were stimulated with DMXAA at 37 °C or 20 °C for the indicated times. Cell lysates were prepared, and EGFP-STING was immunoprecipitated with anti-GFP antibody. The cell lysates and the immunoprecipitated proteins were analyzed by western blotting. **b**, *Sting^-/-^* MEFs stably expressing mRuby3-STING and mNeonGreen (mNG)-Ubiquitin were imaged by Airyscan super-resolution microscopy every 5 min after DMXAA stimulation. Selected frames from Supplementary Video 3 are shown. Scale bar, 10 μm. **c**, Quantitation of the number of mNG-Ubiquitin puncta (Supplementary Video 3). **d**, *Sting^-/-^* MEFs reconstituted with EGFP-STING were treated with control siRNA or *Tsg101* siRNA. The cells were incubated with DMXAA for the indicated times and analyzed by western blotting. **e**, *Sting^-/-^* MEFs reconstituted with mRuby3-STING were treated with control siRNA or *Tsg101* siRNA for 60 h, and then stimulated with DMXAA for 12 h. Cells were fixed, permeabilized, and immunostained with anti-polyubiquitin antibody (cyan) and anti-Lamp1 (yellow). Scale bars, 10 μm. **f, g**, Fluorescence intensity profiles along the red lines in (**e**) are shown. **h**, EGFP-Tsg101 (WT or ΔUEV) and mRuby3-STING were stably expressed in *Sting^-/-^* MEFs. Cells were then treated with DMXAA for the indicated times, fixed, and imaged. Scale bars, 10 μm. **i, j**, Fluorescence intensity profiles along the red lines (**h**) are shown. **k**, siRNA-resistant EGFP-Tsg101 (WT or ΔUEV) and mRuby3-STING were stably expressed in *Sting^-/-^* MEFs. Cells were treated with control siRNA or *Tsg101* siRNA for 60 h, and then stimulated with DMXAA for 12 h. The expression of Cxcl10 was quantitated with qRT-PCR. **l**, A graphical abstract illustrating lysosomal vesiculophagy.

We sought to examine the dynamics of ubiquitin with STING stimulation. *Sting^-/-^* MEFs were stably transduced with mRuby3-STING and mNeonGreen-ubiquitin and imaged with Airyscan super-resolution microscopy. As with EGFP-STING^6^, mRuby3-STING translocated to the perinuclear Golgi by 30 min after DMXAA stimulation (Extended Data Fig. 7a, b), and then to REs by 120 min (Extended Data Fig. 7c). mNeonGreen-ubiquitin distributed diffusively throughout the cytosol and was then translocated to several puncta 120 min after stimulation. These mNeonGreen-ubiquitin puncta were positive with Rab11 (an RE protein) and mRuby3-STING (Extended Data Fig. 7c-e), suggesting that STING underwent ubiquitination at REs. Live cell imaging showed essentially the same results: mNeonGreen-ubiquitin was recruited to mRuby3-STING-positive puncta 95 min after stimulation, at the timing when STING localized at REs (Fig. 5b, c, Supplementary Video 3).

We next examined whether Tsg101, a ubiquitin-binding protein, was required for the degradation of ubiquitinated STING. The smeared bands corresponding to ubiquitinated EGFP-STING diminished 12 hours after stimulation in control cells, but not in cells treated with Tsg101 siRNA (Fig. 5d). In accordance with this biochemical data, the fluorescence of mRuby3-STING and that of polyubiquitin immunostained by anti-polyubiquitin antibody diminished 12 hours after stimulation in control cells, but not in cells treated with Tsg101 siRNA (Fig. 5e-g). Of note, mRuby3-STING and polyubiquitin did not colocalize with lysosomes in Tsg101-depleted cells. These results suggested a role of Tsg101 in encapsulation of ubiquitinated STING into lysosomes.

Finally, we examined a role of an *N*-terminal ubiquitin E2 variant (UEV) domain of Tsg101 in the termination of the STING signalling. EGFP-Tsg101 distributed diffusively throughout the cytosol before stimulation and was translocated to several mRuby3-STING-positive puncta 2 hours after stimulation (Fig. 5h-j). In contrast, a Tsg101 variant lacking the UEV domain remained diffusive and did not translocate to the mRuby3-STING-positive puncta. These results suggested that Tsg101 bound to ubiquitinated STING through its UEV domain. Knockdown of Tsg101 enhanced DMXAA-dependent induction of Cxcl10 (Fig. 3e, Fig. 5k). This enhanced transcription of Cxcl10 was suppressed by the expression of siRNA-resistant form of wild-type Tsg101, but not Tsg101 (ΔUEV). mRuby3-STING lingered in Tsg101-depleted cells 12 hours after stimulation (Fig. 4a). This duration of the fluorescence was suppressed by the expression of siRNA-resistant form of wild-type Tsg101, but not Tsg101 (ΔUEV) (Extended Data Fig. 8a, b). These results indicated that the binding of the UEV domain of Tsg101 to ubiquitinated STING was essential for lysosomal degradation of STING and the termination of the STING signalling.

## Discussion

The degradation of STING at lysosomes is pivotal to prevent the persistent transcription of innate immune and inflammatory genes^5^. Requisites of primary structure of STING for its degradation have been identified, such as phosphorylation of Ser365^5^ and the helix consisting of amino acid residues 281-297^23^. In contrast, the cellular machinery underlying the degradation of STING has not been clear. The stimulation of STING triggers macroautophagy^5,25^, but this appears not to be involved in the degradation of STING^25^ (Extended Data Fig.1a). In the present study, we showed that STING vesicles in the cytosol were encapsulated into lysosomal lumen and degraded there (Fig. 5l). The STING encapsulation was not mediated through fusion of the membrane of STING vesicles with that of lysosome. Thus, this phenomenon can be defined as “lysosomal microautophagy”, by which degradative substrates in the cytosol reach lysosomal lumen directly.

As to lysosomal microautophagy, CMA has been characterized^14,32^. CMA mediates the selective degradation of cytosolic proteins in lysosome. Mechanistically, KFERQ-like motifs present in the substrate proteins are recognized by a cytosolic chaperone protein Hsc70c and directed to Lamp2A at lysosomal surface, followed by the translocation of the substrate proteins through lysosomal membrane. Of note, CMA is not expected to mediate the degradation of transmembrane proteins. In line with this, we confirmed that knockdown of Lamp2 did not interfere with the encapsulation of STING into lysosomes or the expression of stimulation-dependent transcription of Cxcl10 (Extended Data Fig. 8d). We also found that STING stimulation did not cause the noticeable degradation of Gapdh, an authentic substrate of CMA (Extended Data Fig. 8e). Thus, the phenomenon we revealed in the present study may represent a novel mode of lysosomal microautophagy, by which a transmembrane protein in membrane vesicles, not soluble cytosolic proteins, is degraded in lysosomes. With a particular emphasis on the findings that the clustered vesicles were directly encapsulated into lysosomal lumen, we coined this “lysosomal vesiculophagy”.

By screening of mammalian *Vps* genes, we identified several ESCRT proteins as essential regulators of lysosomal vesiculophagy. The ESCRT generates inverse membrane involutions on a variety of organellar membranes and cooperate with the ATPase Vps4 to drive membrane scission or sealing. On early endosomes/late endosomes, the ESCRT plays a key role in the biogenesis of intraluminal vesicles (ILVs), which are destined for degradation at lysosomes or for extracellular secretion as exosomes. Intriguingly, the size of the endosomal ILVs is rather small, ranging between 40-80 nm^33^, which highly contrasts with that of lysosomal ILVs (indicated by the arrows in moss green in Fig. 2b-d), ranging between 200-300 nm. Therefore, the nature of the ESCRT operating on lysosomes may be distinct from that on endosomes. In this regard, it is noteworthy that a component of ESCRT-0, Hgs (also known as Hrs) was dispensable for lysosomal vesiculophagy (Extended Data Fig. 5a).

During or after STING stay at REs, STING was located on a cluster of uniform membrane vesicles with electron-dense coat (Fig. 2e, f). Around the same timing, STING underwent extensive ubiquitination (Fig. 5), and this ubiquitination appeared essential for lysosomal encapsulation of STING. The vesiculation of STING membrane, through the reduction of its size, may facilitate the process of lysosomal encapsulation. Coat proteins may endow STING membranes with stiffness so that lysosomal encapsulation would proceed efficiently.

REs are organelles that function in recycling molecules back to the plasma membrane^10,34^. Besides their classical roles in endocytic recycling, it has been shown that REs have a role in exocytic and retrograde membrane traffic^35,36^, demonstrating that REs serve as a central hub for sorting various cargos to different destinations^37^. In the present study, we revealed that REs also had a role in a previously unanticipated traffic pathway by which an exocytic cargo protein STING was delivered to lysosomes. Given the nature of the pathway that STING follows^38,39^, namely, “ER-Golgi-REs-lysosomes”, lysosomal vesiculophagy may contribute to the proteostasis of exocytic proteins and ER/Golgi resident proteins. It remains to be elucidated whether lysosomal vesiculophagy is a constitutive catabolic process or a catabolic process triggered by certain cellular events, such as the emergence of STING-vesicles with extensive ubiquitination.

## Supporting information

Sup_video_1(TfnR-EGFPmovie)

Sup_video_2(Lysosome live imaging)

Sup_video_3(ubiquitin movie)

## Methods

### Regents

Antibodies used in this study were shown in Supplementary Table 2. The following reagents were purchased from the manufacturers as noted: DMXAA (14617, Cayman), anti-FLAG M2 Affinity Gel (A2220, Sigma), LysoTracker™ Deep Red (L12492, Thermo Fisher Scientific, Waltham, MA), E64d (4321, Peptide Institute), pepstatinA (4397, Peptide Institute), orlistat (O4139, MERCK).

### Cell culture

MEFs were obtained from embryos of WT or *Sting^-/-^* mice at E13.5 and immortalized with SV40 Large T antigen. MEFs were cultured in DMEM supplemented with 10% fetal bovine serum (FBS) and penicillin/streptomycin/glutamine (PSG) in a 5 % CO_2_ incubator. MEFs that stably express tagged proteins were established using retrovirus. Plat-E cells were transfected with pMXs vectors, and the medium that contains the retrovirus was collected. MEFs were incubated with the medium and then selected with puromycin (2 μg ml^-1^), blasticidin (5 μg ml^-1^), or hygromycin (400 μg ml^-1^) for several days. RAW-Lucia™ ISG-KO-STING Cells (InvivoGen) were cultured in DMEM supplemented with 10% fetal bovine serum (FBS), normocin (100 μg ml^-1^), and penicillin/streptomycin/glutamine.

### Immunocytochemistry

Cells were seeded on coverslips (13 mm No.1S, MATSUNAMI), fixed with 4% paraformaldehyde (PFA) in PBS at room temperature for 15 min, and permeabilized with digitonin (50 μg ml^-1^) in PBS at room temperature for 5 min. After blocking with 3% BSA in PBS, cells were incubated with primary antibodies followed by secondary antibodies at room temperature for 1 h. Cells were then mounted with ProLong™ Glass Antifade Mountant (P36982, Thermo Fisher Scientific).

Confocal microscopy was performed using a LSM880 with Airyscan (Zeiss) with a 63 × 1.4 Plan-Apochromat oil immersion lens or 100 × 1.46 alpha-Plan-Apochromat oil immersion lens. Images were analyzed and processed with Zeiss ZEN 2.3 SP1 FP3 (black, 64 bit) (ver. 14.0.21.201) and Fiji (ver. 2.0.0-rc-69/1.52p).

### Live-cell imaging

The day before imaging, cells were seeded on a glass bottom dish (627870, Greiner bio-one). The medium was changed to DMEM^gfp^-2 (MC102, evrogen) containing 10 % FBS, PSG, and rutin (20 μg ml^-1^) (30319-04, nacalai tesque) before imaging. Live-cell imaging was performed using LSM880 with Airyscan (Zeiss) equipped with a 100 × 1.46 alpha-Plan-Apochromat oil immersion lens and Immersol™ 518F/37°C (444970-9010-000, Zeiss). During live-cell imaging, the dish was mounted in a chamber (STXG-WSKMX-SET, TOKAI HIT) to maintain the incubation conditions at 37 °C and 5% CO_2_. Acuired images were Airyscan processed with Zeiss ZEN 2.3 SP1 FP3 (black, 64 bit) (ver. 14.0.21.201) and analyzed with Fiji (ver. 2.0.0-rc-69/1.52p).

### PCR cloning

cDNAs encoding mouse STING, mouse Rab5a, mouse Transferrin receptor (TfnR), human Lamp1, mouse ubiquitin, mouse Tsg101 were amplified by PCR. The cDNAs were inserted into pMX-IPuro or pMX-IBla. Tsg101 [ΔUEV (aa 146-391)] and siRNA-resistant Tsg101 were generated by site-directed mutagenesis.

### Type I interferon bioassay

MEFs were treated with indicated siRNA for 62 h followed by stimulation with DMXAA for 10 h. Cell culture supernatants were then added to RAW-Lucia™ ISG-KO-STING Cells (Invivogen). Twelve hours after incubation, the luciferase activity was measured by GloMax® Navigator Microplate Luminometer (Promega).

### Flow cytometry

*Sting^-/-^* MEFs reconstituted with mRuby3-STING were treated with indicated siRNA for 54 h followed by stimulation with or without DMXAA for 18 h. Cells were detached with trypsin/EDTA and fixed with 4% PFA in PBS at room temperature for 15 min. Mean Fluorescence Intensity was analyzed by Cell Sorter SH800 (Sony).

### qRT-PCR

Total RNA was extracted from cells using ISOGEN II (Nippongene) or SuperPrep II (TOYOBO), and reverse-transcribed using ReverTraAce qPCR RT Master Mix with gDNA Remover (TOYOBO). Quantitative real-time PCR (qRT-PCR) was performed using KOD SYBR qPCR (TOYOBO) and LightCycler 96 (Roche). Target gene expression was normalized on the basis of Gapdh content.

### Immunoprecipitation

Cells were washed with ice-cold PBS and scraped in immunoprecipitation buffer composed of 50 mM HEPES-NaOH (pH 7.2), 150 mM NaCl, 5 mM EDTA, 1% Triton X-100, protease inhibitor cocktail (25955, dilution 1:100) (Nacalai Tesque) and phosphatase inhibitors (8 mM NaF, 12 mM b-glycerophosphate, 1 mM Na_3_VO_4_, 1.2 mM Na_2_MoO_4_, 5 mM cantharidin, 2 mM imidazole). The cell lysates were centrifuged at 15,000 rpm for 15 min at 4 °C, and the resultant supernatants were pre-cleared with Ig-Accept Protein G (Nacalai Tesque) at 4 °C for 15 min. The lysates were then incubated for 3 h at 4 °C with anti-GFP (3E6) and Ig-Accept Protein G. The beads were washed four times with immunoprecipitation wash buffer [50mM HEPES-NaOH (pH 7.2), 150 mM NaCl, 0.1% Triton X-100] and eluted with 2 × Laemmli sample buffer. The immunoprecipitated proteins were separated with SDS-PAGE and transferred to PVDF membrane, then analyzed by western blotting.

### Western blotting

Proteins were separated in polyacrylamide gel and then transferred to polyvinylidene difluoride membranes (Millipore). These membranes were incubated with primary antibodies, followed by secondary antibodies conjugated to peroxidase. The proteins were visualized by enhanced chemiluminescence using Fusion SOLO.7S.EDGE (Vilber-Lourmat).

### Mass spectrometry

Cells were lysed with IP buffer (50 mM HEPES-NaOH (pH 7.2), 150 mM NaCl, 5 mM EDTA, 1% Triton X-100, protease inhibitors, and phosphatase inhibitors). The lysates were centrifugated at 15,000 rpm for 10 min at 4 °C, the resultant supernatants were incubated for overnight at 4 °C with anti-FLAG M2 Affinity Gel. The beads were washed four times with immunoprecipitation wash buffer (50 mM HEPES-NaOH (pH 7.2), 150 mM NaCl, 1% Triton X-100), and eluted with elution buffer [50 mM HEPES-NaOH (pH 7.2), 150 mM NaCl, 5 mM EDTA, 1% Triton X-100, 500 μg ml^-1^ FLAG peptide]. Eluted proteins were applied to SDS-PAGE, and the electrophoresis was stopped when the samples were moved to the top of the separation gel. The gel was stained with CBB and the protein bands at the top of separation gel were excised. The proteins were reduced and alkylated with acrylamide, followed by a tryptic digestion in gel (TPCK treated trypsin, Worthington Biochemical Corporation). The digests were separated with a reversed phase nano-spray column (NTCC-360/75-3-105, NIKKYO technos) and then applied to Q Exactiv Hybrid Quadrupole-Orbitrap mass spectrometer (Thermo Scientific). MS and MS/MS data were obtained with TOP10 method. The MS/MS data was searched against NCBI nr database using MASCOT program 2.6 (Matrix Science) and the MS data was quantified using Proteome Discoverer 2.2 (Thermo Scientific).

### RNA interference

siRNA (siGENOME) used in this study was purchased from Dharmacon. Cells were transfected with siRNA (5 nM) using Lipofectamine RNAiMAX (Invitrogen) according to the manufacturer’s instruction. Six hours after transfection, the medium was replaced by DMEM with 10% FBS followed by incubation for 66 h.

### Quantification of the number of mNeonGreen-ubiquitin puncta

Images of mNeonGreen-ubiquitin were thresholded using Yen’s method with Fiji. mNeonGreen-ubiquitin positive puncta were defined using the “Analyze Particles” menu from Fiji on the binary thresholded image.

### CLEM

Cells were cultured on coverslips coated with 150-μm grids (Matsunami Glass Ind., Ltd.). The cells were stimulated with DMXAA (25 μg ml^-1^), protease inhibitors [E64d (30 μg ml^-1^) and pepstatin A (40 μg ml^-1^)], orlistat (20 μg ml^-1^). Cells were fixed with 2% paraformaldehyde–2% glutaraldehyde in 0.1 M phosphate buffer (pH 7.4) for 15 min at room temperature and rinsed three times for 15 min each time in 0.1 M phosphate buffer (pH 7.4). The fluorescence images were obtained using a confocal microscope [LSM880 with Airyscan (Zeiss)]. They were fixed again with 2% paraformaldehyde-2% glutaraldehyde in 0.1 M phosphate buffer (pH 7.4) for more than 15 min at 4°C, and then with a reduced osmium fixative. After embedding in Epon812 resin, areas containing cells of interest were trimmed according to the light-microscopic observations, and serial ultrathin sections (80 nm-thickness) were prepared and observed with an electron microscope (JEM1400EX; JEOL)^40,41^.

### Statistical analyses

Error bars displayed throughout this study represent s.e.m. unless otherwise indicated and were calculated from triplicate or quadruplicate samples. The data were statistically analyzed by performing Student’s unpaired t-test with Bonferroni multiple correction.

### Data availability

The data sets generated during and/or analyzed during the current study are available from the corresponding author on reasonable request.

## Acknowledgements

This work was supported by JSPS KAKENHI Grant Numbers JP19H00974 (T.T.), JP17H06164 (H.A.), JP17H06418 (H.A.), JP20H03415 (S.W.), JP20H05307 (K.M.), JP20H03202 (K.M.), JP17K15445 (K.M.), and JP21K07104 (T.U.); AMED-PRIME (17939604) (T.T.); JSPS Research Fellowship for Young Scientists (19J21426, Y.K.; 19J23315, E.O.); the Subsidy for Interdisciplinary Study and Research concerning COVID-19 (Mitsubishi Foundation) (T.T.), Takeda Science Foundation (S.W. and K.M.),. the Research Foundation For Pharmaceutical Sciences (K.M.), JST Center of Innovation program from Japan (JPM-JCE1303) (K.M.) Grant for Basic Science Research Projects from the Sumitomo Foundation (K.M.), Koyanagi-Foundation (K.M.), and the Nakatomi Foundation (K.M.). We thank Takayuki Yabe and Katsuyuki Kanno for their technical supports in the EM.

## Author contribution

Y.K. and K.M. designed and performed the experiments, analyzed the data, interpreted the results, and wrote the paper; Y.T. performed siRNA screening; A.S. performed live cell imaging; S.H. performed qPCR analysis; R.U. performed the experiments for identification of STING binding proteins; E.O. designed and performed experiments; T.S. and N.D. performed the proteomics analysis; T.U. performed the experiments with electron microscopy; H.A. designed the experiments, interpreted the results; S.W. performed the experiments with electron microscopy and interpreted the results; T.T. designed the experiments, interpreted the results, and wrote the paper.

## Competing interests

The authors declare no competing financial interests.

---

**Extended Data Fig. 1.**
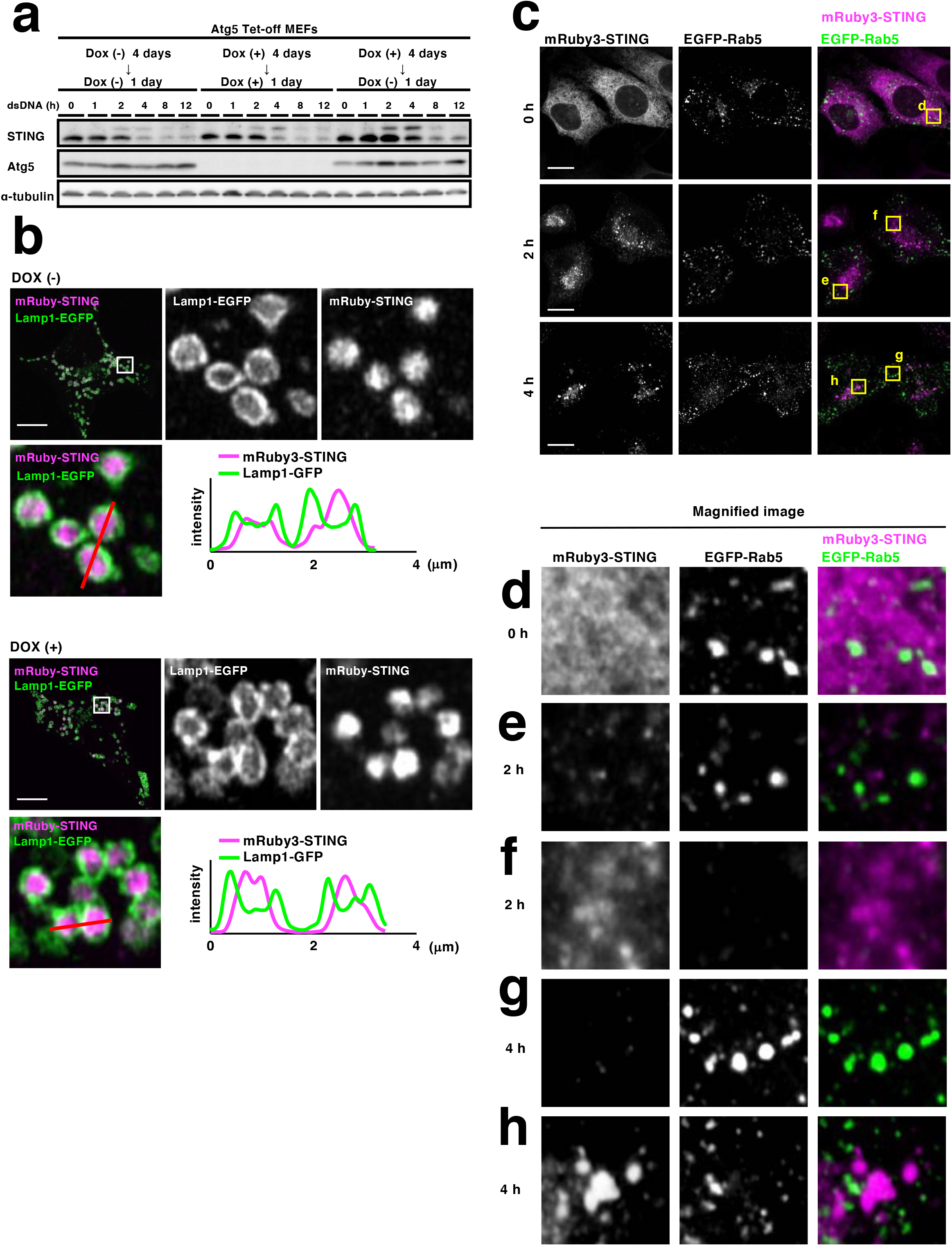
**(related to Figure 1) a**. Atg5 Tet-off cells were cultured with or without Doxycycline (Dox) (10 ng ml^-1^). Cells were stimulated with dsDNA for the indicated times. Cell lysates were then prepared and analyzed by western blotting. **b**, mRuby3-STING (magenta) and Lamp1-EGFP (green) were stably expressed in Atg5 Tet-off cells. Cells were cultured with or without Dox (10 ng ml^-1^) for 4 days. The cells were then treated with DMXAA (25 *μ*g ml^-1^) and protease inhibitors [E64d (30 *μ*g ml^-1^) and pepstatin A (40 *μ*g ml^-1^)] for 12 h. Cells were fixed and imaged by Airyscan super-resolution microscopy. The boxed areas are magnified and shown. Fluorescence intensity profiles along the red lines are shown. Scale bars, 10 *μ*m. **c**, EGFP-Rab5a (green) and mRuby3-STING (magenta) were stably expressed *Sting^-/-^* MEFs. Cells were treated with DMXAA for the indicated times. Cells were fixed and imaged by Airyscan superresolution microscopy. **d-h**. The yellow boxed areas in (**c**) are magnified and shown.

**Extended Data Fig. 2.**
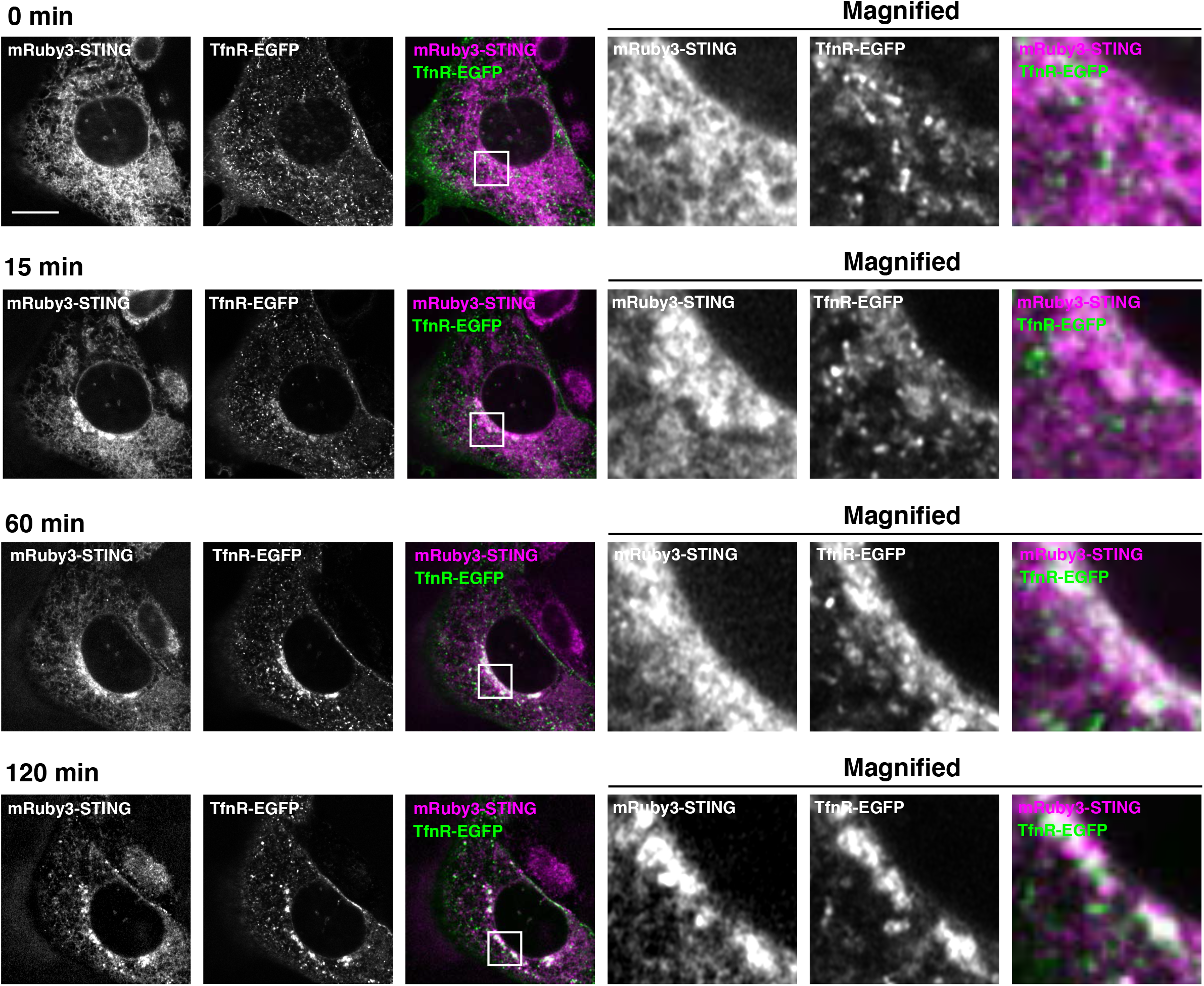
**(related to Figure 1)** *Sting^-/-^* MEFs stably expressing mRuby3-STING and TfnR-EGFP were imaged by Airyscan super-resolution microscopy every 1 min after stimulation with DMXAA (25 *μ*g ml^-1^). Selected images from the movie are shown. See also Supplementary Movie 2. The boxed areas in the merged images are magnified and shown. Scale bar, 10 *μ*m.

**Extended Data Fig. 3.**
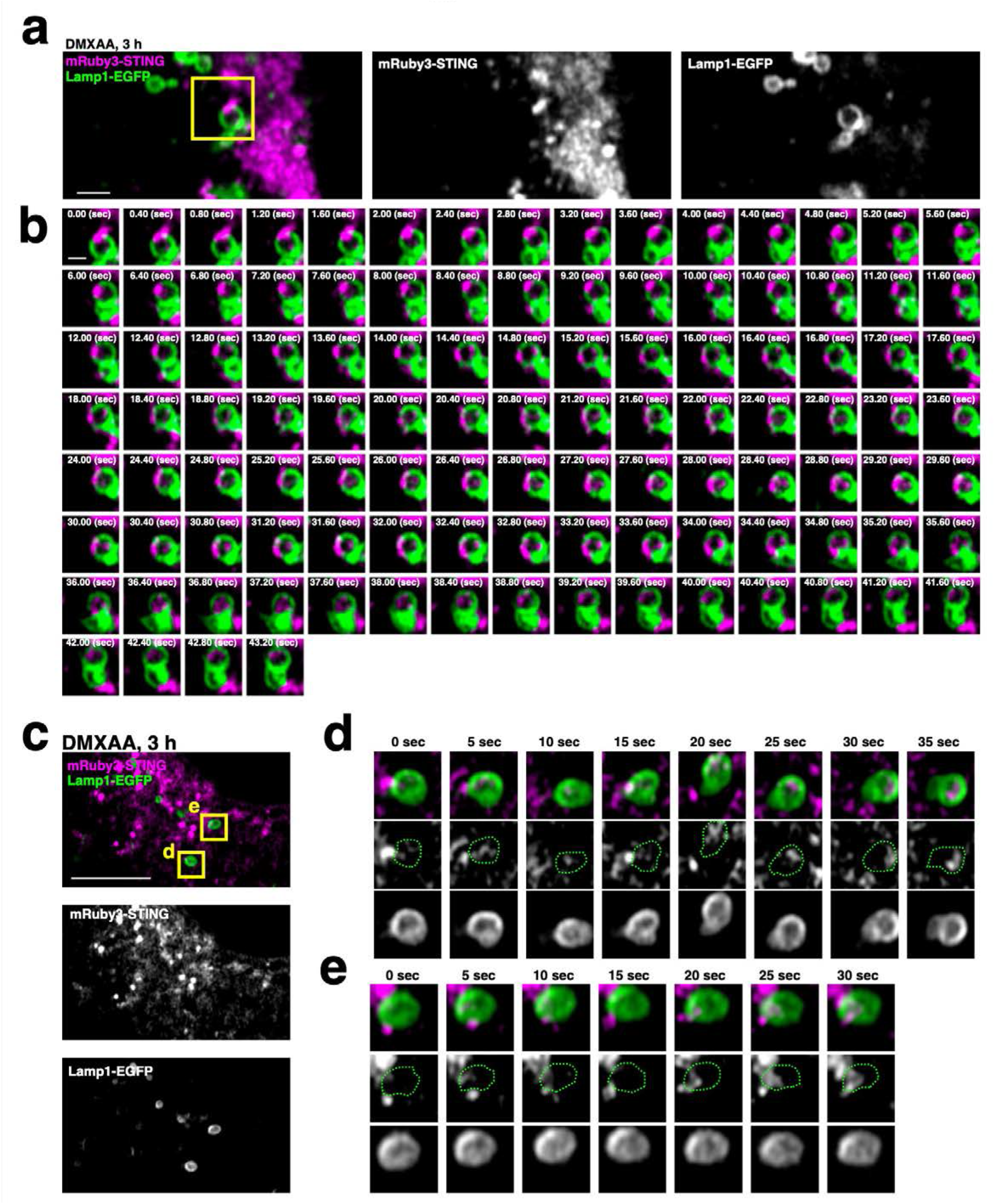
**(related to Figure 1) a,** *Sting^-/-^* MEFs stably expressing mRuby3-STING and Lampi-EGFP were imaged by Airyscan super-resolution microscopy every 0.4 seconds from 3 hours after DMXAA stimulation (related to Fig.1g). **b**, The time-lapse images of the region outlined by the yellow box in (**a**) are shown sequentially. Scale bar, 500 nm. **c-e**, Cells were imaged by Airyscan super-resolution microscopy every 5 seconds from 3 hours after DMXAA stimulation. The time-lapse images of the regions outlined by the yellow boxes in (**c**) are shown sequentially in (**d**) and (**e**). The dotted green lines indicate the limiting membrane of lysosome. Scale bar in (**c**), 10 *μ*m.

**Extended Data Fig. 4.**
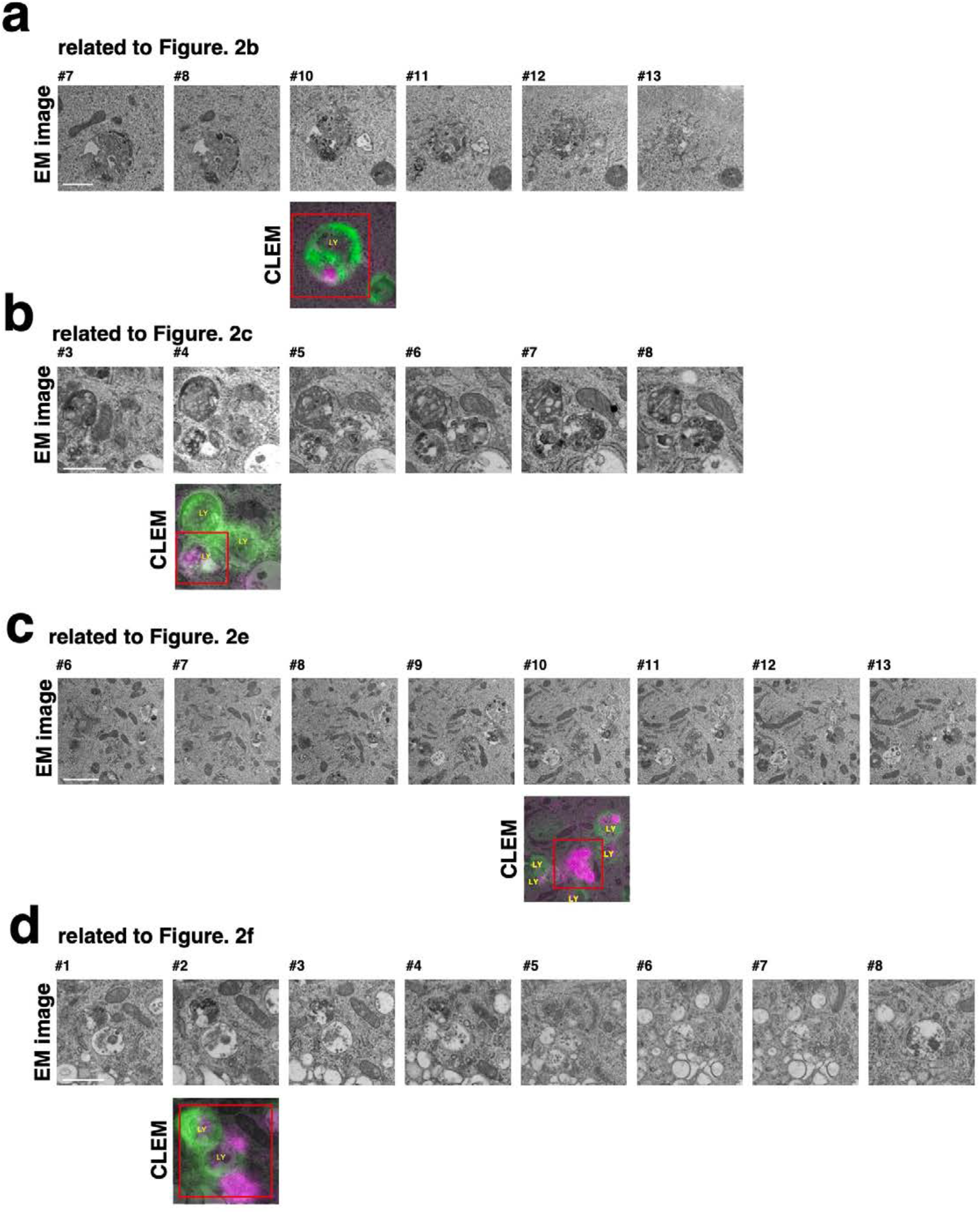
**(related to Figure 2)** Serial EM pictures and a selected CLEM image are shown. Panels **a, b, c,** and **d** correspond to Figs. 2b, 2c, 2e, and 2f, respectively. The number (#) indicates the order in the serial section. LY, Lysosomes; Scale, 500 nm.

**Extended Fig. 5.**
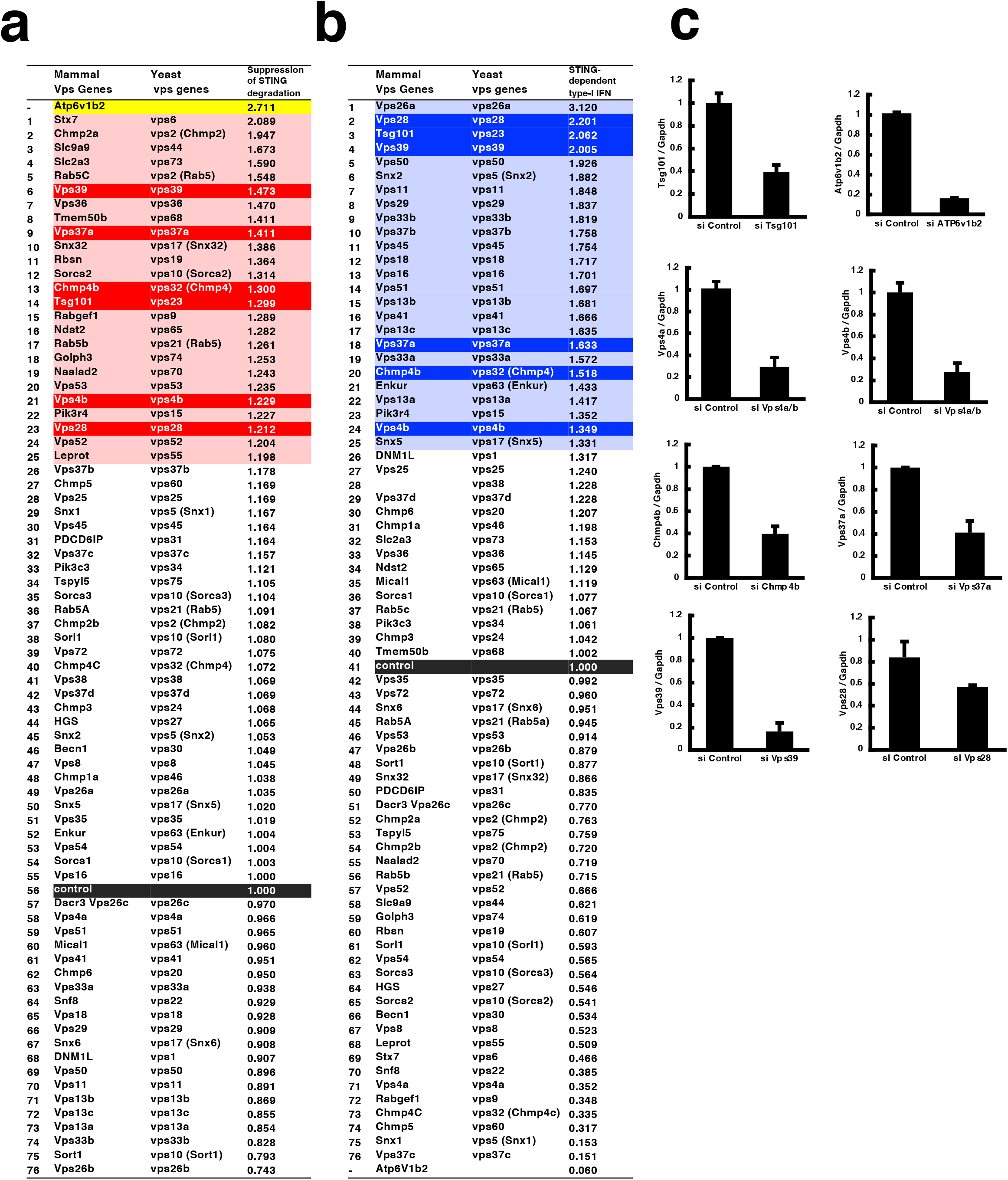
**(related to Figure 3) a**, The data related to Fig. 3b are shown. The top 25 genes are highlighted in red. The genes that were also ranked within top 25 in “type I interferon assay” are highlighted in bright red. **b**, The data related to Fig. 3c are shown. The top 25 genes are highlighted in blue. The genes that were also ranked within top 25 in “STING degradation assay” are highlighted in bright blue. **c**, Knockdown efficiency of Vps genes and Atp6v1b2. Cells were treated with the indicated siRNAs for 72 hours, and qRT-PCR was performed. Gapdh was used as an internal control.

**Extended Data Fig. 6.**
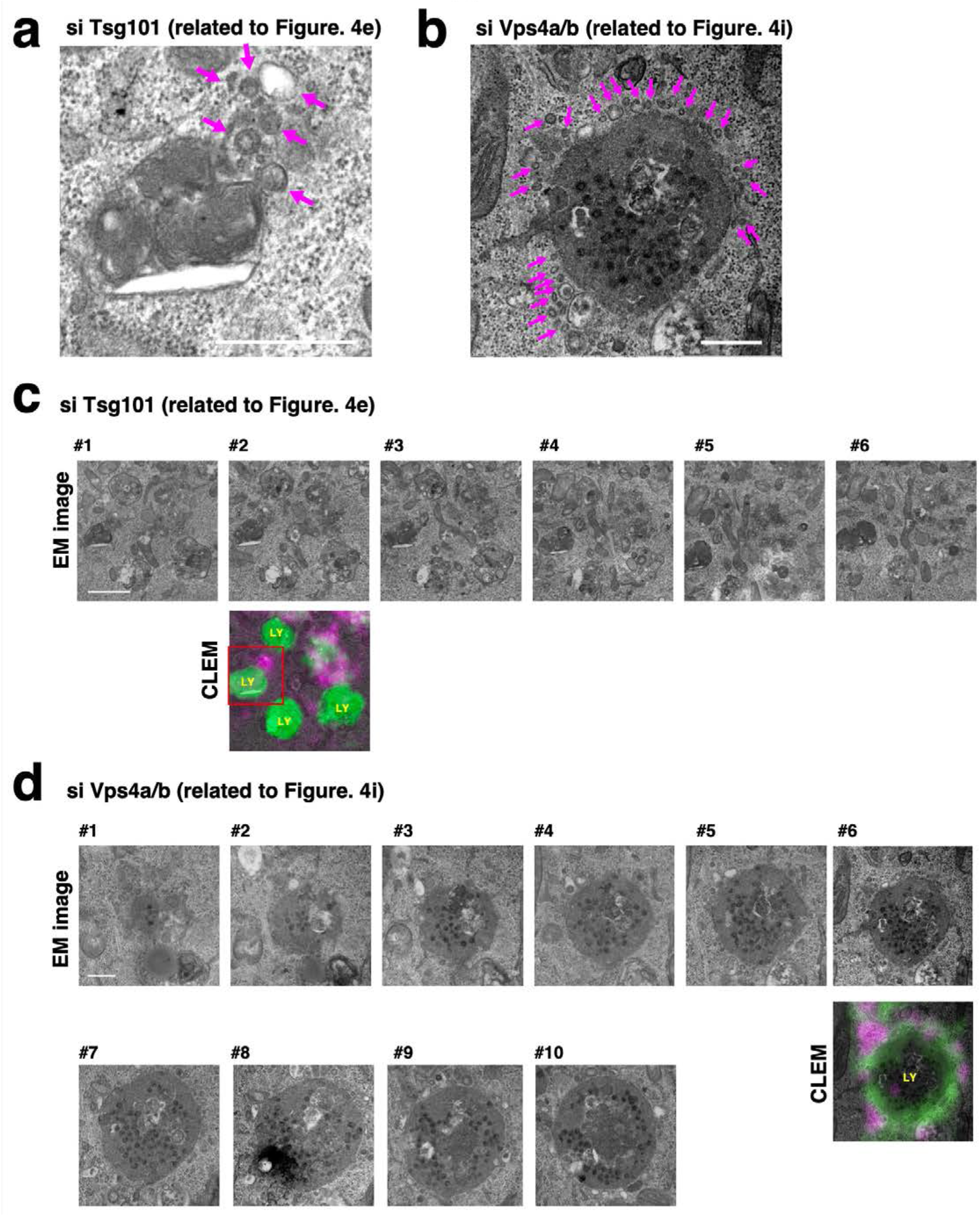
**(related to Figure 4) a,** The magnified EM picture of Fig. 4e. **b,** The magnified EM picture of Fig. 4i. **c** and **d,** Serial EM pictures and a selected CLEM image are shown. Panels **(c)** and **(d)** correspond to Figs. 4e and 4i, respectively. The number (#) indicates the order in the serial section. LY, Lysosomes; Bar, 500 nm.

**Extended Fig. 7.**
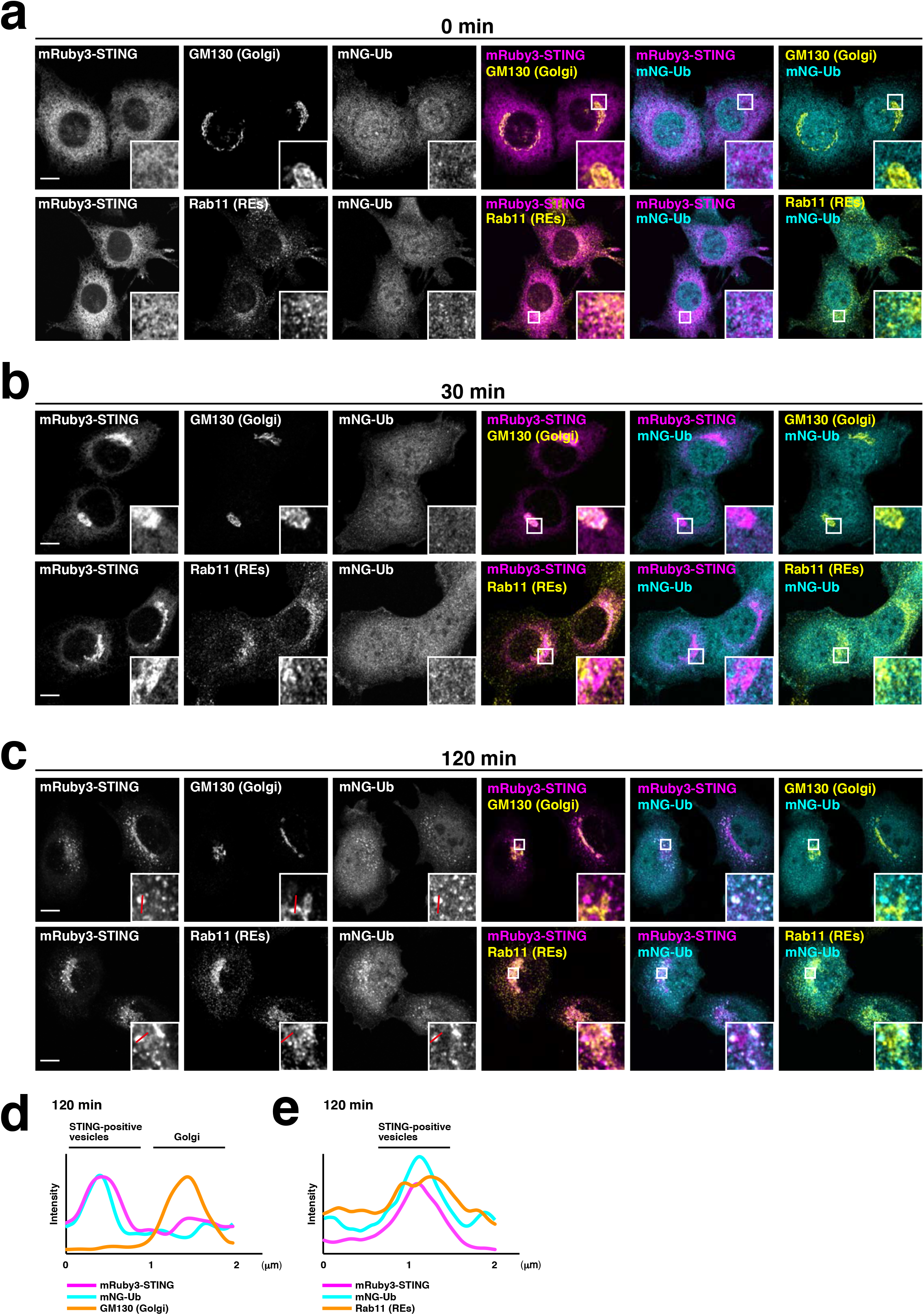
**(related to Figure 5) a-c**, mNeonGreen-ubiquitin (mNG-Ub, cyan) and mRuby3-STING (magenta) were stably expressed in *Sting^-/-^* MEFs. Cells were stimulated with DMXAA for the indicated times. Cells were then fixed, permeabilized, and immunostained with anti-GM130 (a Golgi protein, yellow) or anti-Rab11 (a recycling endosomal protein, yellow) antibodies. **d**, **e**, Fluorescence intensity profiles along the red lines in (**c**) are shown. Scale bars, 10 *μ*m.

**Extended Fig. 8.**
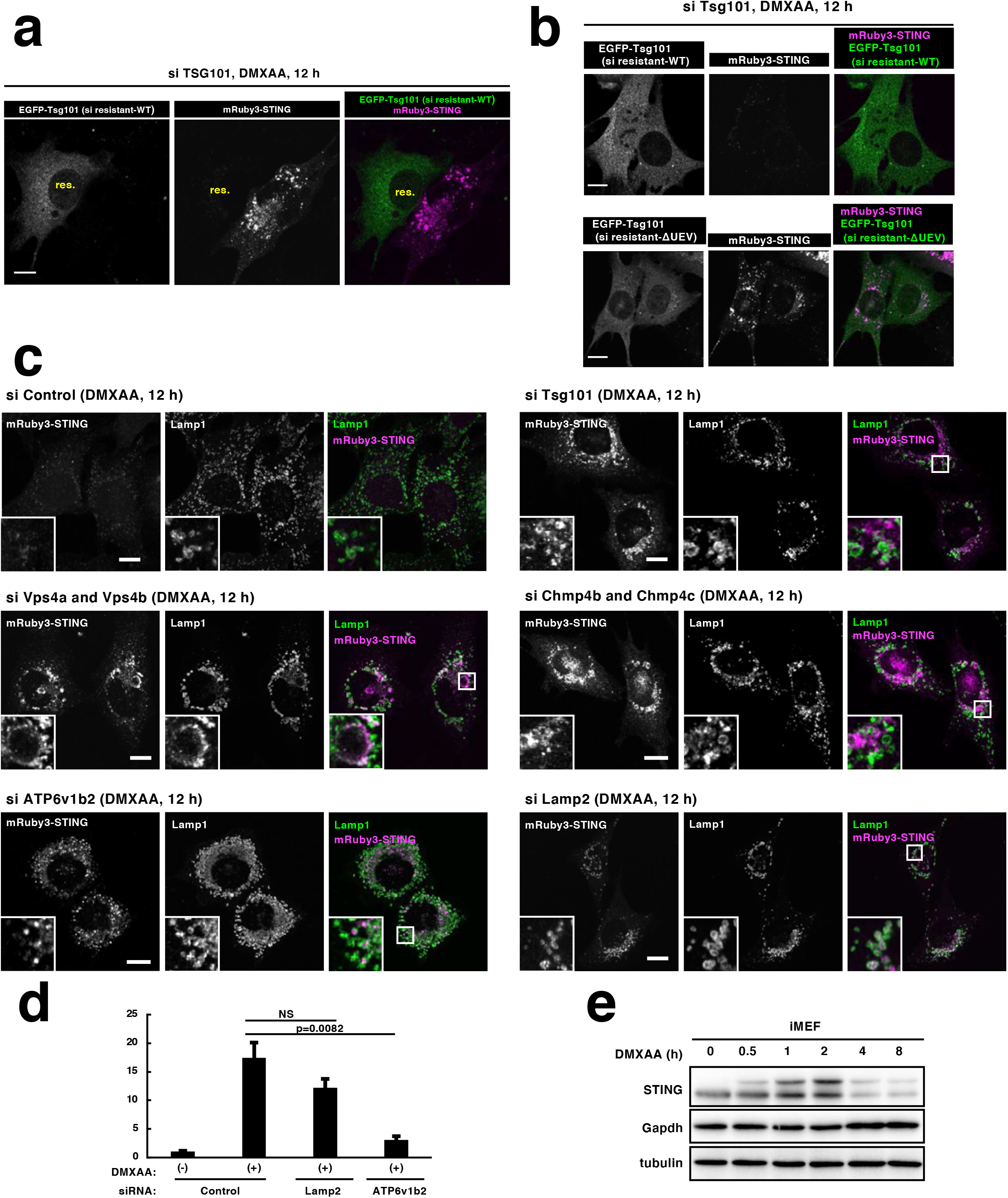
**(related to Fig.5) a**, Two cell lines *Sting^-/-^* MEFs expressing mRuby3-STING, and *Sting^-/-^* MEFs expressing mRuby3-STING and siRNA resistant EGFP-Tsg101 (WT)] were mixed, treated with Tsg101 siRNA for 60 h, and stimulated with DMXAA for 12 h. These cells were then fixed and imaged by Airyscan super-resolution microscopy. res., cells expressing EGFP-Tsg101. **b**, siRNA-resistant EGFP-Tsg101 (WT or ΔUEV) and mRuby3-STING were stably expressed in *Sting^-/-^* MEFs. Cells were treated with Tsg101 siRNA for 60 h and stimulated with DMXAA for 12 h. Cells were then fixed and imaged by Airyscan super-resolution microscopy. **c**, *Sting^-/-^* MEFs expressing mRuby3-STING were treated with the indicated siRNAs for 60 h and stimulated with DMXAA for 12 h. Cells were then fixed, permeabilized, and immunostained with anti-Lamp1 (a lysosomal protein). Scale bars, 10 μm. **d**, qRT-PCR of the expression of Cxcl10 in MEFs that were treated with the indicated siRNAs for 60 h, and then stimulated with DMXAA for 12 h. **e**, Western blots of cell lysates of MEFs stimulated with DMXAA for the indicated times.

**Extended Data Fig.9.**
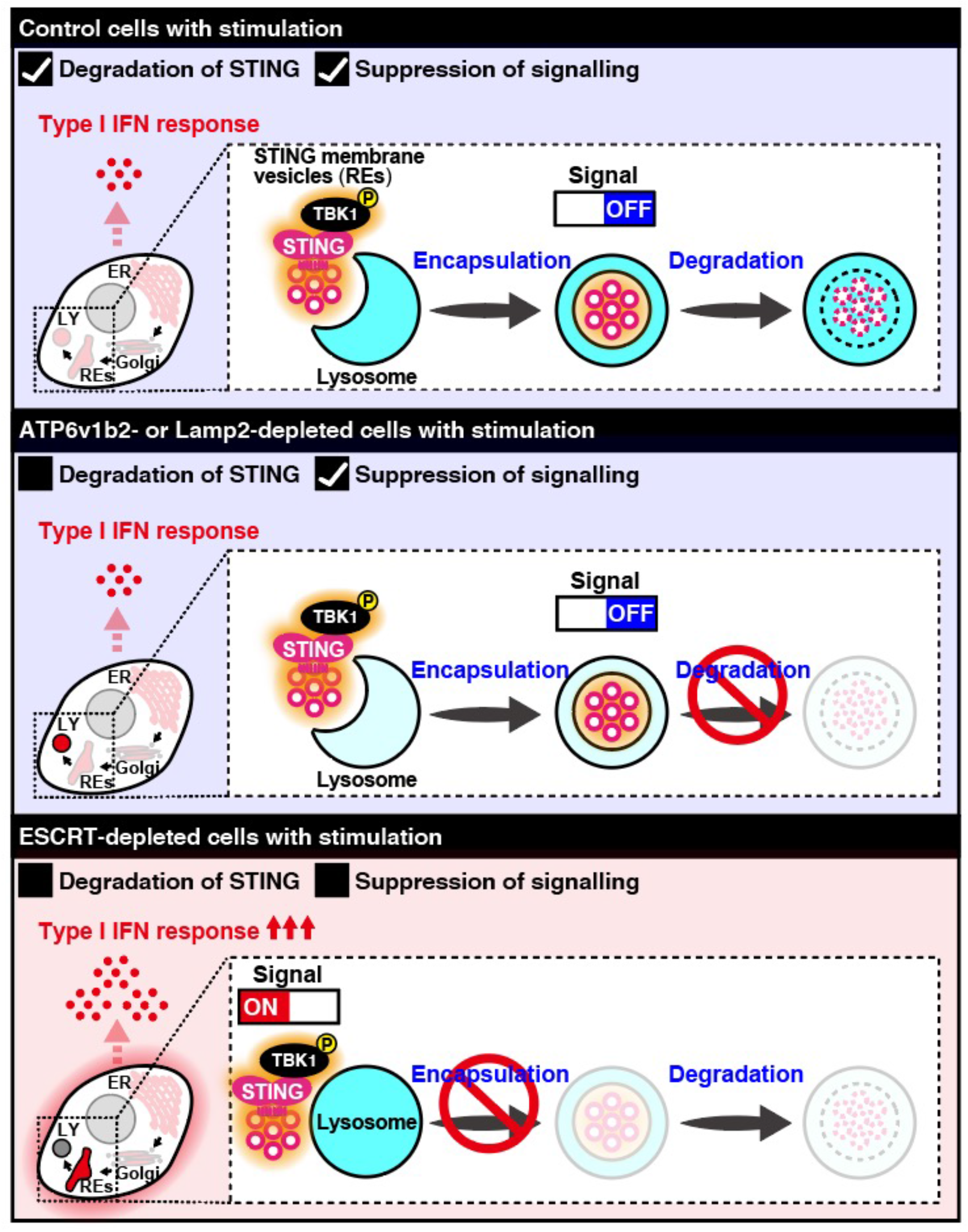
**Graphical abstract of the present study.** (**Control cells**) Active STING/TBK1 complex is encapsulated into lysosomal lumen by “lysosomal vesiculophagy” and eventually degraded there. Thus the signalling is terminated. (**ATP6v1b2- or Lamp2-depleted cells**) The encapsulation of active STING/TBK1 complex proceeds (Extended Fig. 8c). Thus the signalling is terminated (Extended Fig. 8d). Please be noted that STING degradation is impaired because of the defect in the ability of lysosomal proteolysis. (**ESCRT-depleted cells**) The encapsulation is impaired, thus active STING/TBK1 complex remains in the cytosol, leading to the duration of the signalling. Recent studies indicate the involvement of C9orf72 (PMID: 32814898), BLOC1(PMID: 29033128), NPC1(PMID: 34290407) in the STING degradation. The exact site of action of these proteins in the membrane traffic that STING follows, namely, “ER-the Golgi-REs-lysosomes” remains to be elucidated.

## Notes

### Competing Interest Statement

The authors have declared no competing interest.

## References

1. Ishikawa, H. & Barber, G. N. STING is an endoplasmic reticulum adaptor that facilitates innate immune signalling. Nature 455, 674–678 (2008).

2. Barber, G. N. STING: infection, inflammation and cancer. Nat Rev Immunol 15, 760–770 (2015).

3. Ishikawa, H., Ma, Z. & Barber, G. N. STING regulates intracellular DNA-mediated, type I interferon-dependent innate immunity. Nature 461, 788–792 (2009).

4. Saitoh, T. et al. Atg9a controls dsDNA-driven dynamic translocation of STING and the innate immune response. Proc Natl Acad Sci U S A 106, 20842–20846 (2009).

5. Konno, H., Konno, K. & Barber, G. N. Cyclic dinucleotides trigger ULK1 (ATG1) phosphorylation of STING to prevent sustained innate immune signaling. Cell 155, 688–698 (2013).

6. Mukai, K. et al. Activation of STING requires palmitoylation at the Golgi. Nat Commun 7, 11932 (2016).

7. Hopfner, K. P. & Hornung, V. Molecular mechanisms and cellular functions of cGAS-STING signalling. Nat Rev Mol Cell Biol 21, 501–521 (2020).

8. Raymond, C. K., Howald-Stevenson, I., Vater, C. A. & Stevens, T. H. Morphological classification of the yeast vacuolar protein sorting mutants: evidence for a prevacuolar compartment in class E vps mutants. Mol Biol Cell 3, 1389–1402 (1992).

9. Raiborg, C. & Stenmark, H. The ESCRT machinery in endosomal sorting of ubiquitylated membrane proteins. Nature 458, 445–452 (2009).

10. Mellman, I. Endocytosis and molecular sorting. Annu Rev Cell Dev Biol 12, 575–625 (1996).

11. Luzio, J. P., Pryor, P. R. & Bright, N. A. Lysosomes: fusion and function. Nat Rev Mol Cell Biol 8, 622–632 (2007).

12. De Duve, C. & Wattiaux, R. Functions of lysosomes. Annu Rev Physiol 28, 435–492 (1966).

13. Mizushima, N., Levine, B., Cuervo, A. M. & Klionsky, D. J. Autophagy fights disease through cellular self-digestion. Nature 451, 1069–1075 (2008).

14. Kaushik, S. & Cuervo, A. M. The coming of age of chaperone-mediated autophagy. Nat Rev Mol Cell Biol 19, 365–381 (2018).

15. Schuck, S. Microautophagy - distinct molecular mechanisms handle cargoes of many sizes. J Cell Sci 133, jcs246322 (2020).

16. Mijaljica, D., Prescott, M. & Devenish, R. J. Microautophagy in mammalian cells: Revisiting a 40-year-old conundrum. Autophagy 7, 673–682 (2014).

17. Wu, J. et al. Cyclic GMP-AMP is an endogenous second messenger in innate immune signaling by cytosolic DNA. Science 339, 826–830 (2013).

18. Sun, L., Wu, J., Du, F., Chen, X. & Chen, Z. J. Cyclic GMP-AMP synthase is a cytosolic DNA sensor that activates the type I interferon pathway. Science 339, 786–791 (2013).

19. Ogawa, E., Mukai, K., Saito, K., Arai, H. & Taguchi, T. The binding of TBK1 to STING requires exocytic membrane traffic from the ER. Biochem Biophys Res Commun 503, 138–145 (2018).

20. Mukai, K. et al. Homeostatic regulation of STING by retrograde membrane traffic to the ER. Nat Commun 12, 61 (2021).

21. Hu, M. M. et al. Sumoylation Promotes the Stability of the DNA Sensor cGAS and the Adaptor STING to Regulate the Kinetics of Response to DNA Virus. Immunity (2016).

22. Gaidt, M. M. et al. The DNA Inflammasome in Human Myeloid Cells Is Initiated by a STING-Cell Death Program Upstream of NLRP3. Cell (2017).

23. Gonugunta, V. K. et al. Trafficking-Mediated STING Degradation Requires Sorting to Acidified Endolysosomes and Can Be Targeted to Enhance Anti-tumor Response. Cell Rep 21, 3234–3242 (2017).

24. Prabakaran, T. et al. Attenuation of cGAS-STING signaling is mediated by a p62/SQSTM1-dependent autophagy pathway activated by TBK1. EMBO J (2018).

25. Gui, X. et al. Autophagy induction via STING trafficking is a primordial function of the cGAS pathway. Nature 567, 262–266 (2019).

26. Hosokawa, N., Hara, Y. & Mizushima, N. Generation of cell lines with tetracycline-regulated autophagy and a role for autophagy in controlling cell size. FEBS Lett 580, 2623–2629 (2006).

27. Katayama, H., Yamamoto, A., Mizushima, N., Yoshimori, T. & Miyawaki, A. GFP-like proteins stably accumulate in lysosomes. Cell Struct Funct 33, 1–12 (2008).

28. Banta, L. M., Robinson, J. S., Klionsky, D. J. & Emr, S. D. Organelle assembly in yeast: characterization of yeast mutants defective in vacuolar biogenesis and protein sorting. J Cell Biol 107, 1369–1383 (1988).

29. Sancho, E. et al. Role of UEV-1, an inactive variant of the E2 ubiquitin-conjugating enzymes, in in vitro differentiation and cell cycle behavior of HT-29-M6 intestinal mucosecretory cells. Mol Cell Biol 18, 576–589 (1998).

30. Pornillos, O. et al. Structure and functional interactions of the Tsg101 UEV domain. EMBO J 21, 2397–2406 (2002).

31. Griffiths, G. & Simons, K. The trans Golgi network: sorting at the exit site of the Golgi complex. Science 234, 438–443 (1986).

32. Cuervo, A. M. & Dice, J. F. A receptor for the selective uptake and degradation of proteins by lysosomes. Science 273, 501–503 (1996).

33. Edgar, J. R., Eden, E. R. & Futter, C. E. Hrs- and CD63-dependent competing mechanisms make different sized endosomal intraluminal vesicles. Traffic 15, 197–211 (2014).

34. Maxfield, F. R. & McGraw, T. E. Endocytic recycling. Nat Rev Mol Cell Biol 5, 121–132 (2004).

35. Ang, A. L. et al. Recycling endosomes can serve as intermediates during transport from the Golgi to the plasma membrane of MDCK cells. J Cell Biol 167, 531–543 (2004).

36. Uchida, Y. et al. Intracellular phosphatidylserine is essential for retrograde membrane traffic through endosomes. Proc Natl Acad Sci U S A 108, 15846–15851 (2011).

37. Taguchi, T. Emerging roles of recycling endosomes. J Biochem 153, 505–510 (2013).

38. Taguchi, T. & Mukai, K. Innate immunity signalling and membrane trafficking. Curr Opin Cell Biol 59, 1–7 (2019).

39. Taguchi, T., Mukai, K., Takaya, E. & Shindo, R. STING Operation at the ER/Golgi Interface. Front Immunol 12, 1629 (2021).

## References for Methods

40. Arai, R. & Waguri, S. Improved Electron Microscopy Fixation Methods for Tracking Autophagy-Associated Membranes in Cultured Mammalian Cells. Methods Mol Biol 1880, 211–221 (2019).

41. Uemura, T. et al. A cluster of thin tubular structures mediates transformation of the endoplasmic reticulum to autophagic isolation membrane. Mol Cell Biol 34, 1695–1706 (2014).

